# Galectin-3 depletion tames pro-tumoural microglia and restrains cancer cells growth

**DOI:** 10.1101/2023.11.13.563707

**Authors:** Luis Cruz Hernández, María Teresa Sánchez Montero, Alberto Rivera-Ramos, Juan García-Revilla, Rocío Talaverón, Marta Mulero-Acevedo, José Luis Venero, Manuel Sarmiento Soto

## Abstract

The glycoprotein Galectin-3 (Gal-3) is a multifunctional molecule that plays a pivotal role in the initiation and progression of various central nervous system diseases, including cancer. Although the involvement of Gal-3 in tumour progression, resistance to treatment and immunosuppression has long been studied in different cancer types, mainly outside the central nervous system, its elevated expression in myeloid and glial cells underscores its profound impact on the brain’s immune response. In this context, microglia and infiltrating macrophages, the predominant non-cancerous cells within the tumour microenvironment, assume critical roles in establishing an immunosuppressive milieu in diverse brain tumours. Through the utilisation of primary cell cultures and immortalised microglial cell lines, we have elucidated the central role of Gal-3 in promoting cancer cell migration, invasion, and an immunosuppressive microglial phenotypic activation. Furthermore, employing two distinct in vivo models encompassing primary (glioblastoma) and secondary brain tumours (breast cancer brain metastasis), our histological and transcriptomic analysis show that Gal-3 depletion triggers a robust pro-inflammatory response within the tumour microenvironment, notably based on interferon-related pathways. Interestingly, this response is prominently observed in tumour-associated microglia and macrophages (TAMs), resulting in the suppression of cancer cells growth.

## 1. INTRODUCTION

The tumour microenvironment (TME) offers a rich spatiotemporal dynamic of cellular components. Apart from malignant cancer cells, there are a complex myriad of tissue resident and infiltrated peripheral immune system cells. This evolving mosaic of multifunctional cells makes the TME a rather complicated topic of study in the field, with great importance to develop novel therapies that may effectively be used in the most challenging cancers to treat.

Breast cancer has been recently placed as the most diagnosed cancer worldwide, with great affinity to colonise distant organs, specially the brain(1). Thus, depending on tumour subtype, breast cancer brain metastases (BCBM) are very common and represent a major health issue with meagre survival rates amongst patients (ca. 12 months)(1). In turn, one of the most common forms of primary brain tumours is glioblastoma (GB), the deadliest types of brain cancer, where patients who undergo treatment have an average survival measured just in months (ca. 15 months)(2). Despite primary brain tumours and the metastatic colonisation of the brain are completely different diseases, they grow under the influence of the unique immune microenvironment found in the central nervous system, which clearly drives the disease progression(1,3).

BCBM and GB share a common hallmark: they are considered immunologically “cold”, owing to their lack of tumour antigens, inefficient T-cell infiltration and an abundant presence of myeloid-derived immune cells with anti-inflammatory activity(4). Amongst the broad plethora of cells that constitute the TME, the most abundant non-cancerous cell type are microglia and infiltrated macrophages. Tumour-associated microglia and macrophages (TAMs) have been described to develop an anti-inflammatory phenotype and ultimately support BCBM and GB growth (5,6).

In that sense, one key component of the immunosuppressive behaviour of microglia and macrophages is Galectin-3 (Gal-3) (7–9). This β-galactoside-binding protein is long known to be a poor prognostic marker in breast cancer (10) and glioblastoma (11). Given the immunomodulatory role of Gal-3, a potential role of this lectin in driving a pro-tumoural TME in brain tumours is worth being considered. Previous works have shown how activated microglia are the major source of Gal-3 within the brain (7,12–14), together with astrocytes (14). Therefore, this study has focused on the role of TAMs-associated Gal-3, abundantly found in BCBM and GB microenvironment, and its potential implication in driving immunosuppression and cancer cells growth.

Our results describe a strong presence of Gal-3 closely associated to TAMs in BCBM and GB human resections, and brain samples from two different *in vivo* models. Blocking Gal-3 *in vitro,* managed to shape microglia activation, supressing their anti-inflammatory phenotype and inhibiting cancer cells migration and invasion. These results were further validated with the *in vivo* models, where we provide compelling evidence supporting a critical role of Gal-3 in driving TAMs-associated immunosuppression within BCBM and GB microenvironment. Lastly, transcriptomic analysis also showed a strong activation of key innate and adaptive immune response pathways within the TME, thus, tempering pro-tumoural TAMs phenotype towards a more pro-inflammatory state, and contributing to cytotoxic T-cell infiltration with the subsequent tumour growth reduction.

## 2. MATERIAL AND METHODS

### Cell culture and transfection

GL261 (mouse glioblastoma), BV2 (mouse microglia) and EO771 (mouse mammary carcinoma) cell lines were purchased from the American Type Culture Collection (ATCC, USA). For detailed culture conditions see Supp. data. Cells were knocked down by small interfering RNA (siRNA) targeting galectin-3 (Santa Cruz Biotechnology, USA ref: sc-155994-SH) according to the manufacturer’s siRNA transfection protocol.

### Migration and invasion studies

Cell migration studies were performed using the scratch wound assay method. For full information please see Supp. data (Supp. Figure 1A). Recombinant galectin-3 (Gal-3) was produced by the Lund-Protein Production Platform (Lund University, Sweden)(15)

For the invasion studies, cell lines were co-cultured using transwell clear inserts with polyester (PET) membrane (Corning, USA ref: 3470). *In vitro* cell invasion for these cell lines was assessed after 24h seeded in a corning BioCoat matrigel invasion chamber assay (Corning, ref: 354480). The experiment and subsequent measurement of cell invasion was performed according to the manufacturer’s Cell Invasion Assay protocol.

### RNA extraction and quantitative PCR

RNA was extracted from the cell lines using the RNeasy Kit (Qiagen, Netherlands). For protocol and primers information see Supp. data.

### Western blotting assay

Membranes were incubated with anti-Gal 3 antibody (1:1000). GAPDH antibody (Sigma-Aldrich; 1:2000; USA) was used as a loading control. For protocol information see Supp. data.

### Animals and surgery

Experiments were performed in 12-week-old mice C57BL/6 and Galectin-3 null mutant mice with the same background, both lines obtained from Charles Rivers. Female mice were used for the BCBM model and male mice for the GB model and. Animal experimentation was carried out in accordance with the European Community Council Directives (86/609/EU) and Spanish law (R.D. 53/2013 BOE 34/11370-420, 2013) for the use and care of laboratory animals. All the animal procedures in this study were previously approved by ethics committee of University of Seville and Junta de Andalucía. Animals were housed under a 12h light/dark cycle with free access to food and water.

Female C57BL/6 mice were injected with 1µl of PBS containing 10^3^ syngeneic mouse mammary carcinoma EO771 cells by unilateral stereotaxic intracerebral injection in the left striatum (16). Male C57BL/6 mice followed the same surgical protocol as above, with 1µl with 10^4^ syngeneic mouse glioblastoma cells GL261(17). 10 and 21 days after injections, animals were culled and brain samples harvested.

### Immunohistochemistry

Staining protocol was performed as previously described (18). For full information regarding protocol and list of antibodies, please see Supp. data.

### Confocal microscopy

Sections were imaged using a Leica Stellaris 8 STED confocal laser scanning microscope with a ×20 objective. Images of wild-type and transgenic mice were taken on the same day and under the same conditions (laser intensities and photomultiplier voltages). Morphometric analysis of fluorescence-labelled structures was performed offline using LAX software and Fiji ImageJ (National Institutes of Health).

### Quantitation of tumour burden

All data were analysed blinded to experimental conditions. Tumour growth was histologically assessed as previously described, (19,20) (Supp. Figure 2). For full information see Supp. data.

### TAMs analysis

Colocalisation studies were analysed using the ImageJ software package (NIH) as previously described (18). Full information on the plugins and software can be found in the Supp. Data.

### Microarray analysis

To study the differences in genes expression between wt (n=3) and Gal-3 KO mice (n=3), Transcriptome Analysis Console (TAC) Software was used. Gene Set Enrichment analysis (GSEA) and the Database for Annotation, Visualization and Integrated Discovery (DAVID) were also used to further analyse gene clusters comparisons and key biological pathways. For full information see Supp. data.

### Statistical analysis

All individual measurements constitute independent biological replicates and the experiments were repeated at least three times (for quantifications, the n is specified in the figure legends). All data are presented as mean ± SD. Unless otherwise stated, data were analysed using GraphPad Prism (v8.4). For 2 groups, Student’s t-test (2-tailed) was used. For more than 2 groups, 1-way ANOVA with Tukey’s multiple comparison test or two-way Anova with Bonferroni test were used and adjusted P values are reported. Statistical significance was defined as p< 0.05.

### Data availability statement

Transcriptomic data were generated at the Genome and Sequencing facilities in the Institute of Biomedicine of Seville (IBiS, Spain). Raw data supporting the findings of this study are completely available from the corresponding author (MSS) on request.

## 3. RESULTS

### TAMs are a major source of Gal-3 in the TME of BCBM and GB

Firstly, we inquired whether the TME of BCBM and GB showed significant levels of Gal-3 and, more importantly, whether it was closely related to TAMs. The *in vivo* EO771 (Figure 1A-C, BCBM) and GL261 (Figure 1D-F, GB) models presented a strong presence of Gal-3 in the TME. Using Iba-1, a pan-marker of TAMs, we observed that most Gal-3 expression colocalised with TAMs. Importantly, our transgenic Gal-3 Knockout (Gal-3KO) model showed no presence of Gal-3, except for a negligible amount likely derived from the tumour cells.

**Figure 1.**
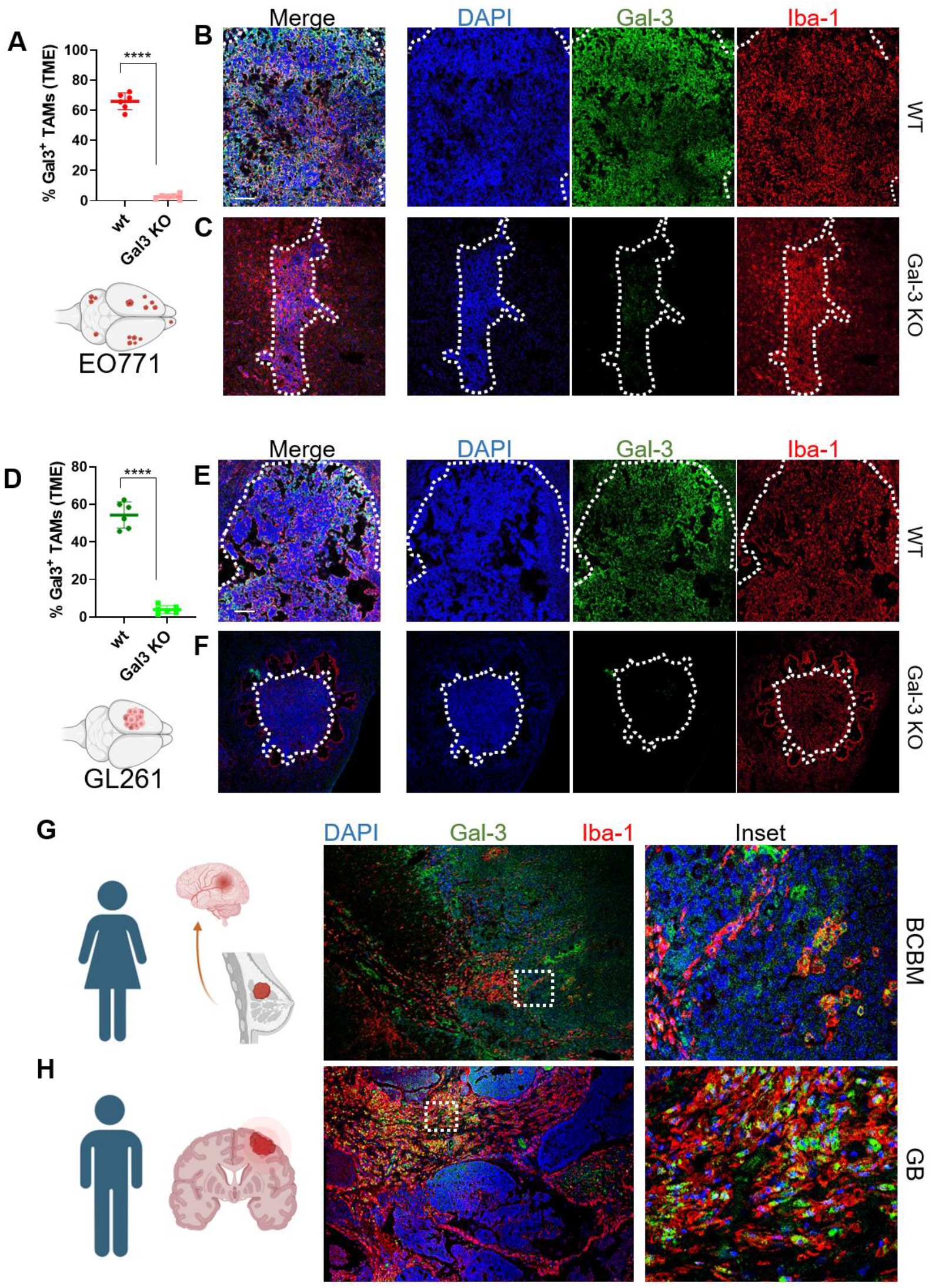
A.-) Quantitation of TAMs expressing Gal-3 within the tumour area in wt and Gal-3KO mice (n=6) injected with EO771 cells. B.-) Imunofuorescence image (merge, left) showing the metastatic area in a wt mouse. (DAPI, blue), Gal-3 (Green) and TAMs (Iba1, red). C.-) Metastatic area of a Gal-3 KO mouse. (Markers as per B). D.-) Quantitation of TAMs expressing Gal-3 within the tumour area in wt and Gal-3KO mice (n=6) injected with GL261. E.-) Imunofuorescence image (merge, left) showing the glioblastoma area in a wt mouse. (DAPI, blue), Gal-3 (Green). and TAMs (Iba1, red). F.-) Metastatic area of a Gal-3 KO mouse. (Markers as per E). G.-) Breast cancer brain metastasis resection from a 64-years-old woman, showing a strong presence of TAMs (Iba1, red) colocalising with Gal-3 (green). H.-) Glioblastoma resection from a 67-years-old man, highlighting (inset) the area with strong presence of TAMs (Iba1, red) and Gal-3 (green). Scale bar: 100µm. *, p<0,0001 (Welch’s t-test).

Finally, human brain resections from patients with BCBM (Figure 1G) and GB (Figure 1H) showed similar patterns, observing expression of Gal-3 in the tumour foci. That Gal-3 expression was again closely related with the presence of TAMs and tumour cells, amongst other cell types. Altogether, with these results, we demonstrated a strong presence of Gal-3-expressing TAMs in two different forms of brain tumours.

### Gal-3 triggers an anti-inflammatory phenotype in microglia cell line

Previous works from our research team described the key role of Gal-3 in TREM2-associated microglial activation (7,12,13). That phenotypic change in microglial response upon Gal-3 presence strongly suggested the need to elucidate the potential role of this lectin in the activating phenotype in TAMs.

Key pro– and anti-inflammatory markers were measured in BV-2 cells upon different tumour-conditioned medium (TCM) challenges. Cells were exposed to TCM from parental tumour cells (EO771 or GL261). Furthermore, in order to study the specific influence of tumoural-derived Gal-3, we silenced Gal-3 in EO771 and GL261 cells and cultured BV2 with the resulting Gal-3-defficient TCM (Gal3KD). Finally, we aimed at elucidating whether exogenous addition of soluble Gal-3 could revert the microglial phenotype found under Gal3KD conditions. Thus, we added soluble Gal-3 back into the medium (+Gal3, Figure 2, Supp. Figure 1). All results were compared with BV-2 coated in normal medium (control condition, DMEM).

**Figure 2.**
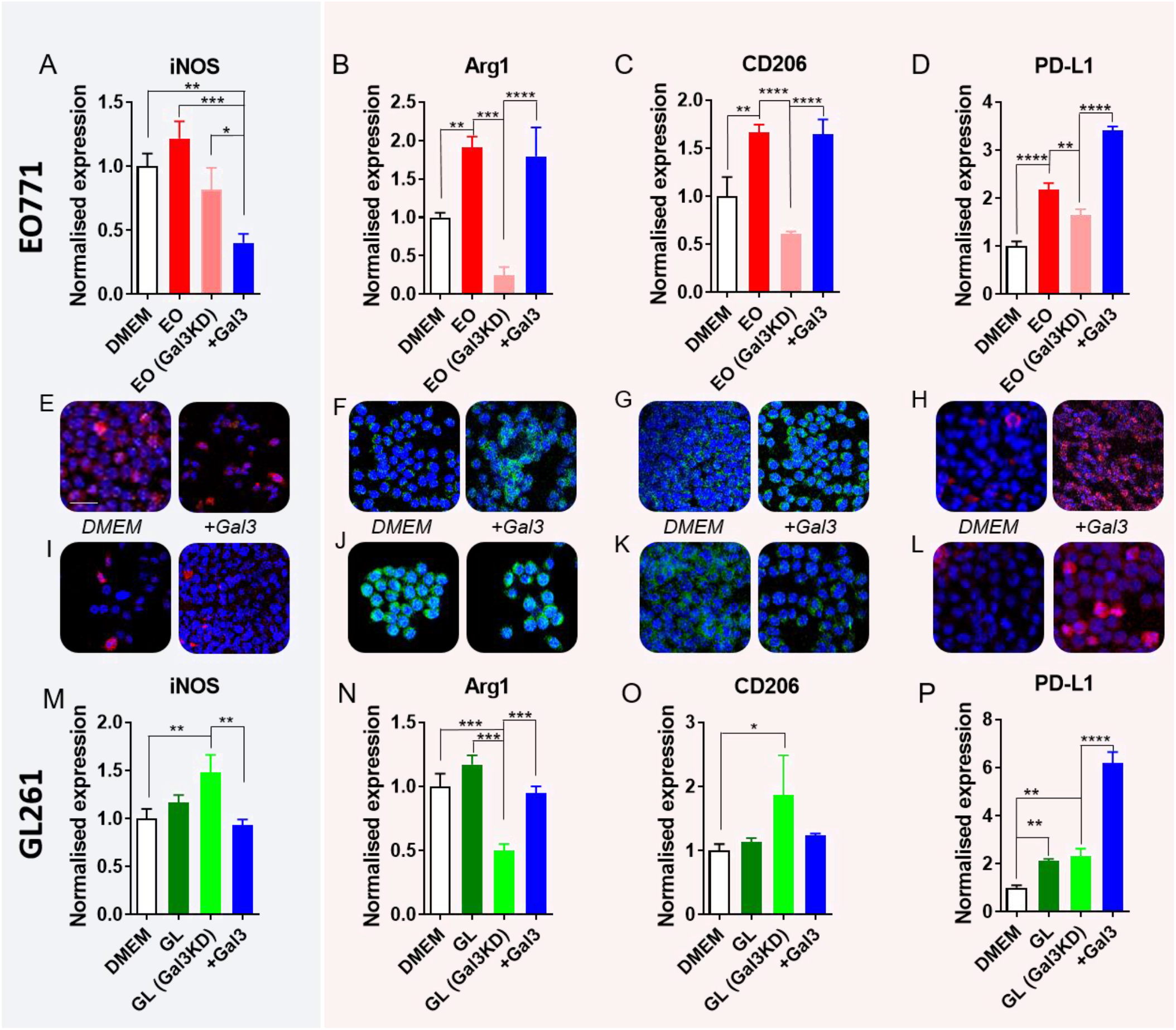
Top row is devoted to BV-2 cells treated with EO771-conditioned medium. Bar graphs show the quantitative analysis of pro-inflammatory marker iNOS (A), and immunosuppressive Arg1 (B), CD206 (C) and PD-L1 (D). Cells were treated (24h) with tumour-conditioned medium from parental EO771 (EO) or EO771 cells with Gal-3 knockdown (siRNA, Gal3KD). Last experimental condition was used to study the effect of the exogenous addition of Gal-3 to EO-Gal3KD medium. Confocal Images showing the expression of iNOS (E, red), Arg1 (Green, F), CD206 (Green, G) and PD-L1 (H, red). Left images refer to control (DMEM) and right images to cells treated with exogenous Gal-3. As per EO771 group, Bottom row is devoted to BV-2 cells treated with GL261-conditioned-medium. Confocal images show expression of iNOS (red, I), Arg1 (Green, J), CD206 (K, Green) and PD-L1 (L, red). Bar graphs show quantitative analysis of the expression of iNOS (M), Arg1 (N), CD206 (O) and PD-L1 (P). Scale bar: 50µm

### EO771-conditioned medium

We aimed at deciphering how tumour-cell-derived Gal-3 may regulate microglia polarisation. To this end, we monitored iNOS, the prototypic marker of pro-inflammatory microglia, and Arg1, CD206 and PD-L1, prototypic markers of anti-inflammatory (pro-tumoural) microglia.

iNOS, a key pro-inflammatory marker (21), (Figure 2A) suffered a significant drop when soluble Gal-3 was added to the medium. Arg1, CD206 and PD-L1 (Figure 2B-D), typical anti-inflammatory markers (6,22,23) had a similar behaviour upon Gal-3 challenges. Once TCM was added, their expression was significantly increased. However, when Gal-3 was silenced in tumour cells and their conditioned medium was then added (Gal3KD), levels of every marker significantly dropped. In contrast to iNOS, when exogenous Gal-3 was added back to the medium, the levels of every immunosuppressive marker were upregulated. Representative images of BV2 cells stained against the above-mentioned markers are depicted in Figure 2E-H.

At a gene level, iNOS showed a similar behaviour as previously described, since its gene expression was significantly upregulated (>3-fold) after EO-TCM and Gal-3 exposition, compared to DMEM. As per seen by immunofluorescence, Arg-1 and PD-L1 were upregulated once BV2 were exposed to TCM and Gal-3. (Supp. Figure 3).

All these data support the role of Gal-3 as a critical promoter of an anti-inflammatory phenotype in microglia, by upregulating classical immune-suppressive markers.

### GL261-conditioned medium

When TCM from Gal-3 KD tumour cells was added (Gal3KD), iNOS was significantly increased in BV-2 (Figure 2M). Interestingly, when Gal-3 was exogenously added, iNOS expression dropped to control levels. Conversely, the absence of Gal-3 in the TCM (Gal3KD) dropped Arg1 levels (Figure 2N), whilst Gal-3 addition had an opposing effect, increasing Arg1 expression. Regarding mannose receptor (CD206), its levels remained unaltered except when cells were coated with Gal3KD-medium (ca. 2-fold increase, Figure 2O). Finally, PD-L1 was always upregulated at any experimental condition (Figure 3P), with special relevance upon soluble Gal-3 addition (ca. 6-fold increase).

**Figure 3.**
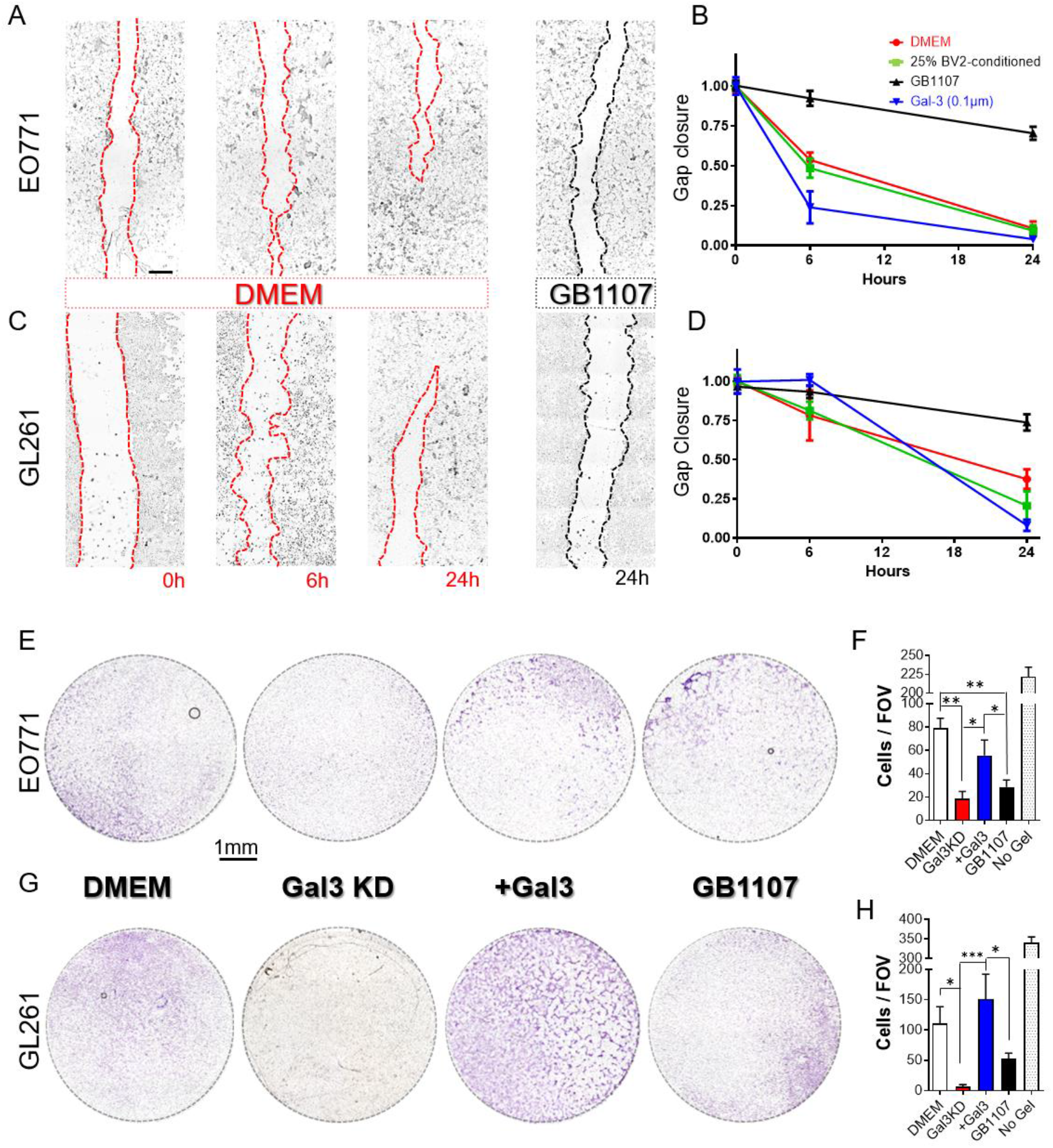
A.-) Scratch-wound assay performed with EO771 cells in DMEM (red dotted lines) conditions. Time-course study with pictures at 0, 6 and 24h. EO771 cells after 24h exposition to GB1107 (black dotted line). B.-) Graph measuring gap closure ability of cells at different experimental conditions: control (DMEM, red line), microglia-conditioned medium (BCV2, Green line), selective Gal-3 inhibitor (GB1107, black line) and after addition of exogenous Gal-3 (blue line). C.-) Pictures of the scratch assay performed to GL261 cells in control (DMEM, red dotted line) and GB1107 condition (black dotted line). D.-) Quantitation of gap closure under the same conditions as described in B. E.-) Invasion assay performed in Boyden Chambers to EO771 cells, stained with cresyl violet. Cells were coated with parental BV2 cells (DMEM) at the bottom well, or BV2 with Gal3 knockdown (siRNA –Gal3KD-). Next experimental condition was performed using exogenous Gal-3 with Gal3KD BV2 cells seeded at the bottom (+Gal3), and finally, using parental BV2 cells with GB1107 (0,1µM). F.-) Quantitation of number of invading cells. G.-Same as E, with GL261 cells. H.-) Same as F with GL261 cells. * p<0,05; ** p<0,01; *** p<0,005. One-way anova with Tukey post hoc analysis. Scale bar 750µm

In the same vein, suggesting the influence of Gal-3 stimulating the immunosuppressive phenotype of BV-2, qPCR analysis showed how Gal-3 increased the gene expression of PD-L1 and Arg-1 even higher than the presence of TCM. In the case of iNOS, both Gal-3 and TCM inhibited the expression of iNOS, thus, suggesting an anti-inflammatory behaviour in BV-2 cells upon Gal-3 presence (Supp. Figure 3)

These data are in agreement with our previous findings regarding EO771-conditioned medium in BV-2 cells, highlighting the key role of Gal-3 upregulating classical immunosuppressive markers (e.g. Arg1 and PD-L1) in BV2 cells.

### Galectin-3 triggers primary microglia activation

Once established the significant role of Gal-3 in BV-2, and recalling that Gal-3 is predominantly expressed and released by activated microglia (7,13,24), we proceeded to further validate Gal-3 impact in primary microglia cell cultures. Pro-inflammatory and tumour supportive phenotypes were characterised upon exposition (24h) to recombinant Gal-3 (+Gal-3). We analysed levels of iNOS and CD80 (25) as classical pro-inflammatory markers, and, on the other side, Arg-1 and PD-L1 as typical tumour-supportive markers (Supp. Figure 4).

*<u>Pro-inflammatory effect</u>*. Wt microglia exposed to soluble Gal-3 induced a significant increase in iNOS and CD80 (Supp. Figure 4A-B). However, in Gal3KO microglia, the addition of Gal-3 had an opposing behaviour, downregulating the expression of iNOS and upregulating CD80.

*<u>Tumour-supportive effect</u>*. Arg-1 (Supp. Figure 4C), the prototypic tumour-supporting marker, was greatly increased once wt microglia was exposed to Gal-3 (<3-fold increase). However, in Gal3KO microglia, Arg1 levels remained unaltered. These results match with the idea that Gal-3 supports immunosuppressive phenotype of immune cells (26,27).

Additionally, PD-L1 (Supp. Figure 4D), a key immune checkpoint inhibitor (4), had a significant increase in any microglial backgrounds after Gal-3 addition. These data match with previous results where Gal-3 upregulated other immune-suppressing markers, further supporting the notion that Gal-3 presence supports a pro-tumoural phenotype in microglia.

### Pharmacological blocking of Galectin-3 hinders tumour cells migration

Considering the potent and multifaceted immunomodulatory functions of Gal-3, we postulated that pharmacological blocking of Gal-3 (GB1107)(28), could be a valid strategy against tumour cells migration. To investigate this, we used EO771 and GL261 to study their migrating properties in response to GB1107 throughout a time-course study.

Interestingly, both cell lines (Figure 3A-D) showed similar sensitivity to Gal-3 modulation. Both tumour cells exposed to microglial (BV2) conditioned medium (Figure 3B and D, green line) had similar gap closure speed across the different time points compared with control (DMEM) conditions (Figure 3A-D, red line). A slight increase in the migration pattern was observed when 0.1µM soluble Gal-3 (Figure 3B-D, blue line) was added to the medium. Importantly, when both tumour cell lines were exposed to GB1107 (selective Gal-3 inhibitor), their migrating properties were greatly reduced (Figure 3A-D, black line). Overall, our results sustain the relevant role of Gal-3 in tumour cell migration.

### Microglial Galectin-3 supports tumour cell invasion

In line with our previous experiments, we aimed at studying the role of microglial Gal-3 in the invading properties of EO771 and GL261 cells. Thus, we designed an invasion assay to study their capacity to spread through an artificial basement membrane (29). Tumour cells were co-cultured with parental BV2 cells, and BV2 cells with Gal-3 expression knocked-down (Gal3 KD) (Supp. Figure 1B). By one side, the number of EO771 cells that successfully reached the luminal side of the membrane was significantly lower with BV2 Gal-3 KD cells at the bottom of the wells (Figure 3E-F and Supp. Figure 1C, Gal3KD). Under these conditions, adding exogenous Gal-3 back into the medium resulted in a noteworthy and significant increase in the number of invading cells (+Gal3). Confirming this notorious role of Gal-3 in supporting tumour cell invasion, the use of GB1107 (0.1µM), mimicked the results from Gal-3 KD condition, and successfully reduced the number of invaded tumour cells (Figure 3E-F, GB1107).

Similarly, GL261 cells mirrored the invasion properties of breast cancer cells when the presence of Gal-3 was modulated (Figure 3G). GL261 cells dropped their invasion capacity (Figure 3G-H) when coated with Gal-3KD BV-2 cells (Gal3KD). Again, when Gal-3 was added back to the medium (+Gal3), the number of cells reached similar values to control conditions (DMEM). Importantly, when GB1107 was added, the number of invasive cells significantly dropped, thus, confirming the important effect of Gal-3 regarding invading capacity of both tumour cell lines (Figure 3H, GB1107).

These results underscore the significant impact of Gal-3 in modulating critical aspects of brain tumours malignancy. Specifically, the potential role of Gal-3 regulating the TME, exerting a robust influence on tumour cells migration and invasion.

Consequently, after all our *in vitro* results, we next proceeded to analyse the effects of Gal-3 in brain tumour growth and TAMs polarisation *in vivo*.

### Glioblastoma and breast cancer brain metastasis burden is reduced in Galectin-3 knockout mice

In order to test *in vivo* the potential impact of the pro-tumoural role of Gal-3, EO771 and GL261 cells were implanted in wild-type C57Bl/6 and Gal-3 KO mice (n=6, per tumour model). Then, the tumour burden was quantified at different time points (Figure 4, 10 and 21 days).

**Figure 4.**
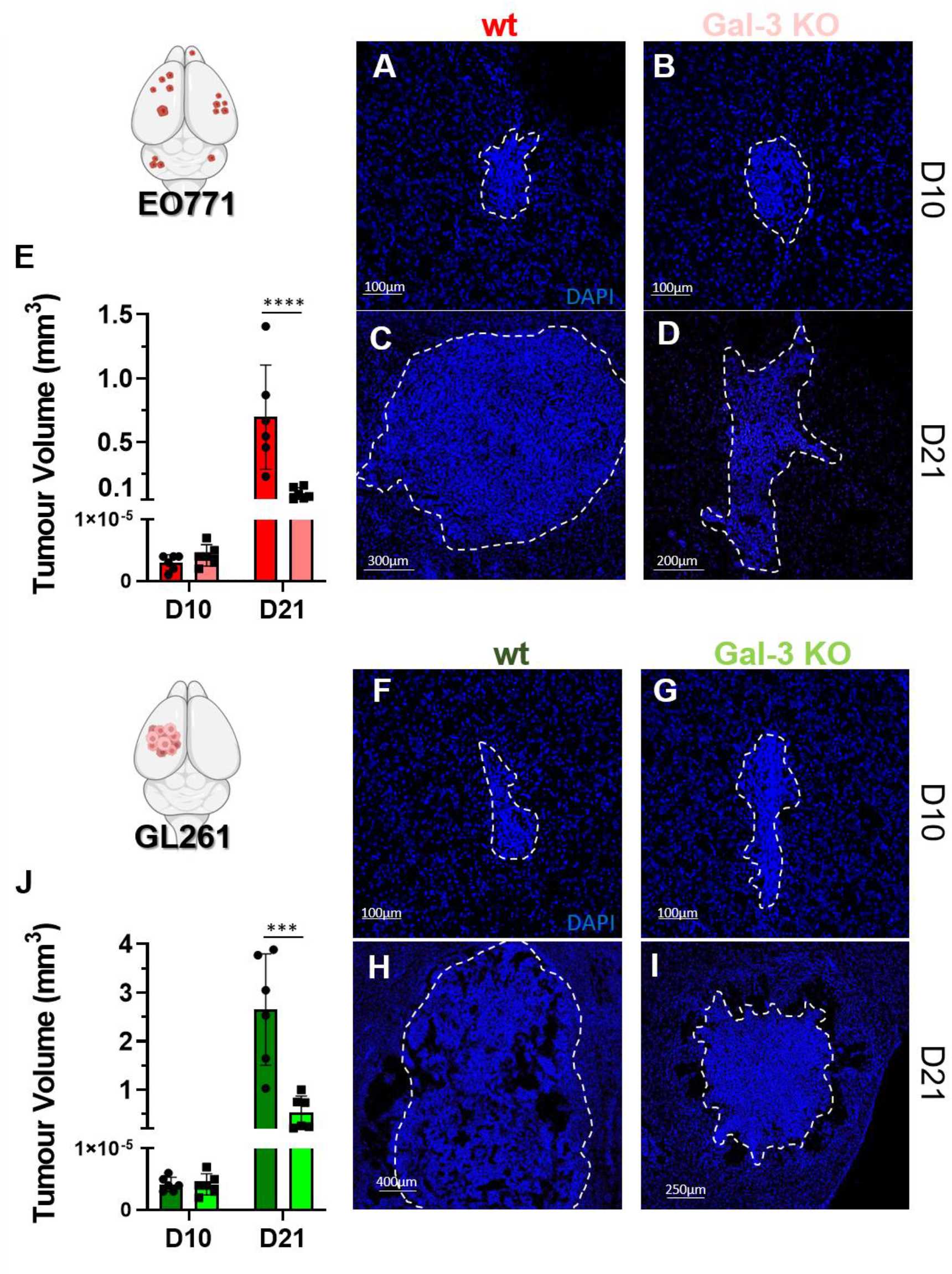
Confocal images of the metastatic area of wt (A) and transgenic Gal3KO (B) female C57Bl/6 at day 10 and day 21 (C and D). E.-) Quantitation of the metastatic area in wt (red) and Gal-3KO mice (light red bars). Confocal images of glioblastoma area in wt (F) and Gal3KO (G) mice injected with GL261 cells at day 10. H and I are images taken at day 21 post injection. J.-) quantitation of glioblastoma area comparing wt (dark Green) and Gal3KO mice (light Green) at 10 and 21 days. Statistical analysis of tumour volume progression was made independently across both times. One-way anova and Tukey post hoc tests was performed. ***, p<0,005. ****, p<0,001.

### Breast cancer brain metastasis

Since 99% of breast cancer cases account for women (30), we focused our BCBM *in vivo* study in female mice. Evaluation of the metastatic burden within the brain of wt and Gal-3 KO C57Bl/6 mice showed no differences at day 10 (Figure 4A-B). However, at day 21, Gal-3KO mice showed a significant drop of the metastatic burden (Figure 4C-E, ca. 8-fold).

### Glioblastoma

Similar to the BCBM model, glioblastoma size showed no difference at day 10 (Figure 4F-G). Again, at day 21, the tumour size was significantly reduced in Gal-3 KO mice (Figure 4H-J, ca. 5-fold).

### Galectin-3 depletion shifted TAMs activation towards a pro-inflammatory phenotype

Using STED confocal microscopy, we quantified immunofluorescent colocalisation of key pro– and anti-inflammatory markers with TAMs. In both models, we clearly observed a phenotype shift of TAMs towards a pro-inflammatory phenotype over immunosuppression (Figure 5).

**Figure 5.**
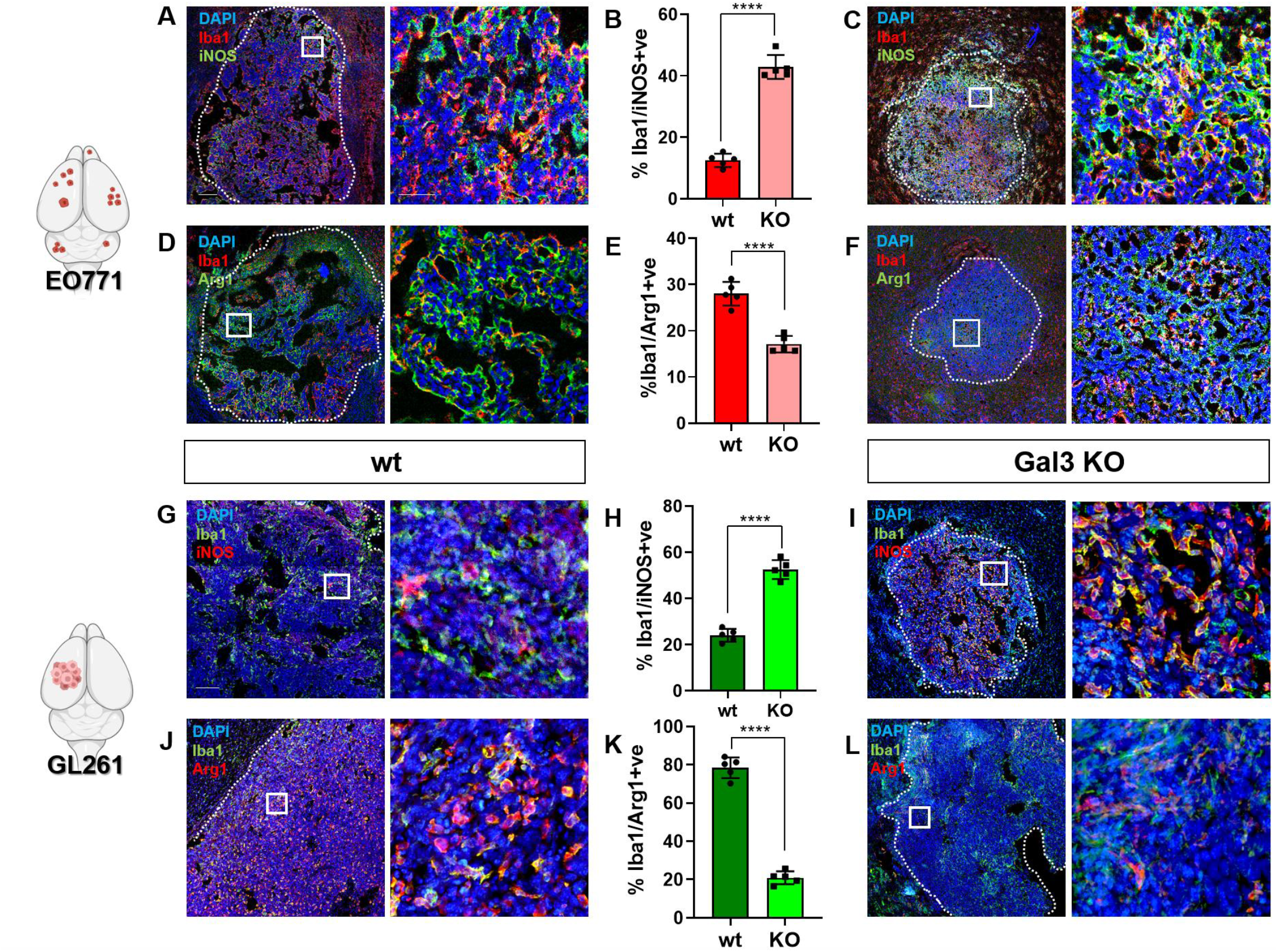
A.-) Confocal images of the metastatic area of wt mouse at day 21 after being injected with EO771 cells. Pictures shows the TME (left image) and the inset (right) with nuceli stained in blue (DAPI), microglia in red (Iba-1) and iNOS (green). B.-) Quantitation of TAMs expressing iNOS within the TME (n=5). C.-) As per A, images showing the TME of Gal3KO mice. D.-Same study as above-mentioned but analysing TAMs expressing Arg1 (E) in wt (D) and Gal3KO (F). G.-) Glioblastoma area of wt mouse injected with GL261 cells at day 21. Pictures shows the TME (left image) and the inset (right) with nuceli stained in blue (DAPI), microglia in green (Iba-1) and iNOS in red. H.-) Quantitation of TAMs expressing iNOS (n=5). I.-) As per G, images showing the TME of Gal3KO mice. J.-) Same study as above-mentioned but analysing TAMs expressing Arg1 (K) in wt (J) and Gal3KO mice (L). Scale bars 250µm (50µm insets).

No differences were found at day 10 after tumour implantation in either model (data not shown). However, at day 21, in the BCBM model, the expression of iNOS within the TAMs population was increased from 15% in wt animals, to 40% in the Gal-3 KO model (Figure 5A-C). Importantly, Arg-1-expressing TAMs population was reduced ̴50% in Gal-3 KO mice compared with wt (Figure 5D-F). Again, these findings suggest the important role of Gal-3 in driving the pro-tumoural TAMs phenotype.

With regard to the GB model, percentage of iNOS-expressing TAMs increased from 23% (wt) up to 58% (Gal-3KO) (Figure 5G-I). On the contrary, the strong immunosuppressive phenotype found in wt mice owing to the high expression of Arg-1, was significantly dropped in Gal-3 KO mice (4-fold decrease, Figure 5J-L).

Understanding the complexity of characterising the TME, owing in great part to the broad number of potential pro– vs anti-inflammatory markers to be used, the iNOS/Arg-1 balance (Supp. Figure 5) may suggest that after Gal-3 depletion, TAMs showed a shift towards an anti-tumoural activation by increasing pro-inflammatory iNOS and reducing the expression of immunosuppressive Arg-1 (17).

### Gal-3 KO mice showed a robust anti-tumour T cell infiltration

Tumour-infiltrating lymphocytes (TILs) are also important components of the TME. High level of CD4^+^ TILs combined with low CD8^+^ TILs is associated with poor prognosis in brain tumours (31). Since Arg-1 expressing TAMs impair T cell responses by modulating the bioavailability of L-arginine and inducing immunosuppression (17), we pursuit the potential effect of Gal-3-dependent TAMs polarisation in TILs population.

In BCBM, the intratumoural CD4^+^ T cells infiltration was significantly increased within the TME of wt mice compared to Gal-3 KO mice (Figure 6A-B). Importantly, CD8^+^ T cells infiltration had an opposing pattern and were more abundant in Gal3 KO mice, again in both tumour models (Figure 6C-D). This CD8^+^/CD4^+^ T Cell ratio has long been described as a good prognosis marker in brain tumours (31).

**Figure 6.**
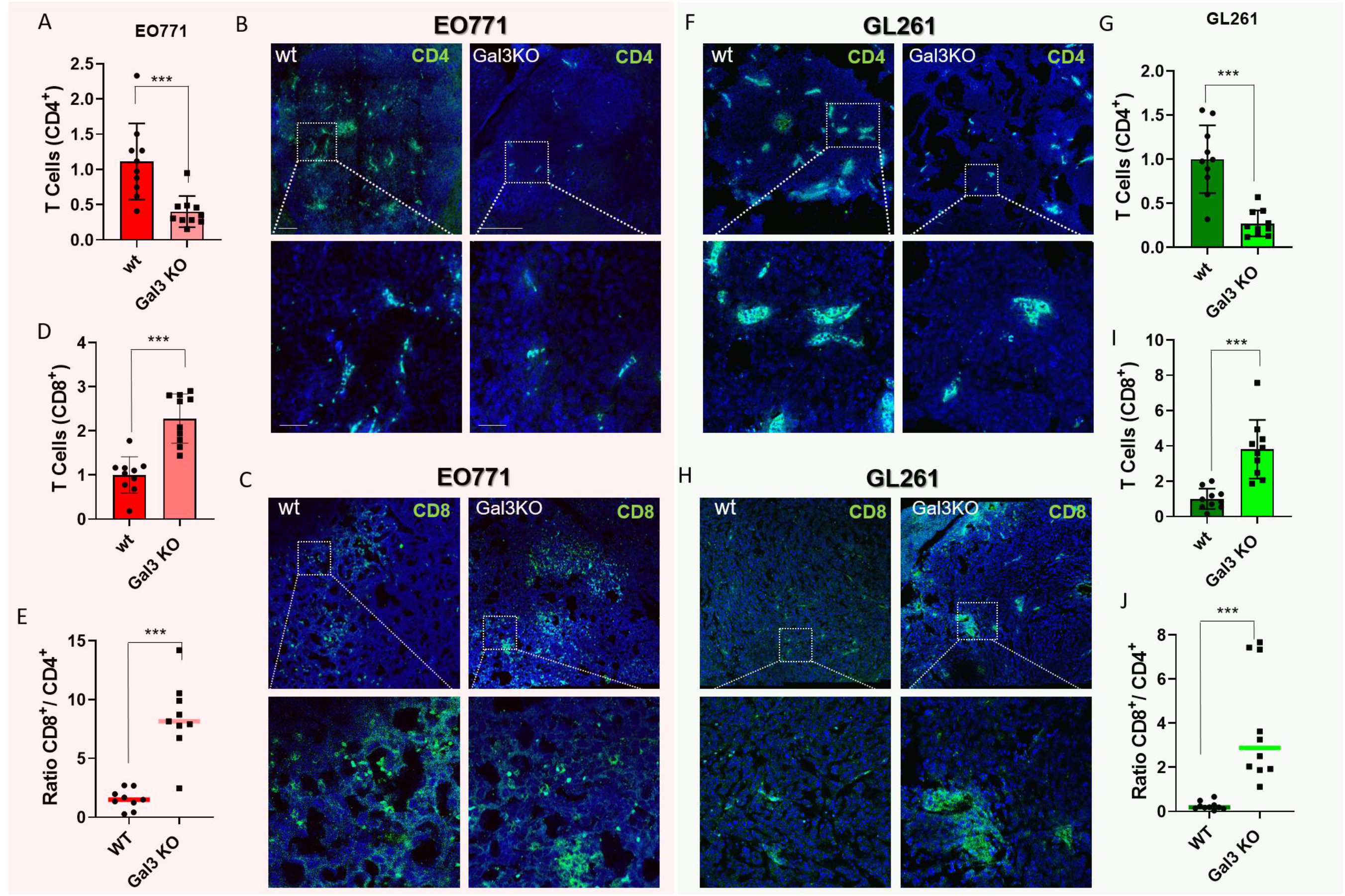
A.-) Quantitation of CD4+ T cells within the TME (green) of BCBM. Nuclei in blue (DAPI) B.-) Confocal images of a BCBM showing CD4+ T cells (green) in the TME in wt and Gal3KO mice. C.-) Confocal images showing CD8+ T cells (green) in the TME of EO771 mice. D.-) Quantitation of CD8+ T cells within the TME of BCBM _1_ mice. E.-) Ratio of CD8 vs CD4 T cells within the TME of BCBM animals. F.-) Confocal images of CD4+ T cells in the TME of GL261-injected mice. G.-) Quantitation of CD4+ T cells (green) in the TME of GB model. H.-) confocal images of CD8+ T cells (green). I.-) Quantitation of intratumoural CD8+ T cells in our GB model J.-) CD8/CD4 T cells ratio within the TME of our GB model. Scale bar 250µm (100µm inset)

In the GB model, T Cells infiltration mirrored the results found in the BCBM mice. CD4^+^ T cells were less infiltrated in the Gal3 KO mice (Figure 6F-G), whilst CD8^+^ T cell population was significantly increased when compared with wt mice (Figure 6H-I).

Interestingly, TAMs, as antigen-presenting cells, have a deep impact in T cell infiltration pattern. Our data suggest that the pro-inflammatory phenotype found in TAMs in Gal3 KO mice, may support a tumouricidal CD8^+^/CD4^+^ TILs ratio (Figure 6E and J).

### Gal-3 depletion enhanced a multifarious pro-inflammatory gene expression signature in the TME

In an initial effort to explore the pre-clinical relevance of our findings, we performed next generation transcriptome-wide gene-level expression profiling studies in the TME. Tumours from BCBM and GB animals were harvested at day 21 post-implantation, and then analysed.

### Breast cancer brain metastasis

Comparing the TME of wt and Gal3 KO mice, we observed 180 dysregulated genes (100 genes upregulated and 80 genes downregulated). The most of upregulated genes in the Gal-3 KO model were immune system-related, closely linked to pro-inflammatory markers (e.g. Tnfsf10, Tnfrsf1b), commonly expressed in microglia and macrophages (e.g. CD68, P2ry14), T cells (e.g. CD2, CD2, CD8b, Granzyme A-B) and Killer cells (Klrb1, KLr1a). On the contrary, wt animals showed pro-metastatic genes expression (e.g. Serpinb9f, Serpinb3b)(32) or classical immunosuppressive markers including arginase-1 (33,34) (Figure 1A). In line with this, gene set enrichment analysis (GSEA) comparing over 22000 genes across both experimental models, showed several pro-inflammatory gene clusters upregulated in the Gal-3 KO model. Specifically, and in accordance with our previous studies, microglial response to interferon gamma was one of the most dysregulated clusters across groups (Figure 7B-C), which is well-known to play a pivotal role in driving the pro-inflammatory phenotype in TAMs (35). Moreover, using wikipathways analysis from TAC software, Gal-3 KO provoked an increase in several gene signatures from different inflammatory responses, with ADAR1 editing deficiency immune response and Type II interferon signalling as the most upregulated pathways (Figure 7D). These pathways are mutually related since ADAR1 is a critical IFN-inducible form, displaying a pivotal role in driving pro-inflammatory responses of the immune system (36,37)

**Figure 7.**
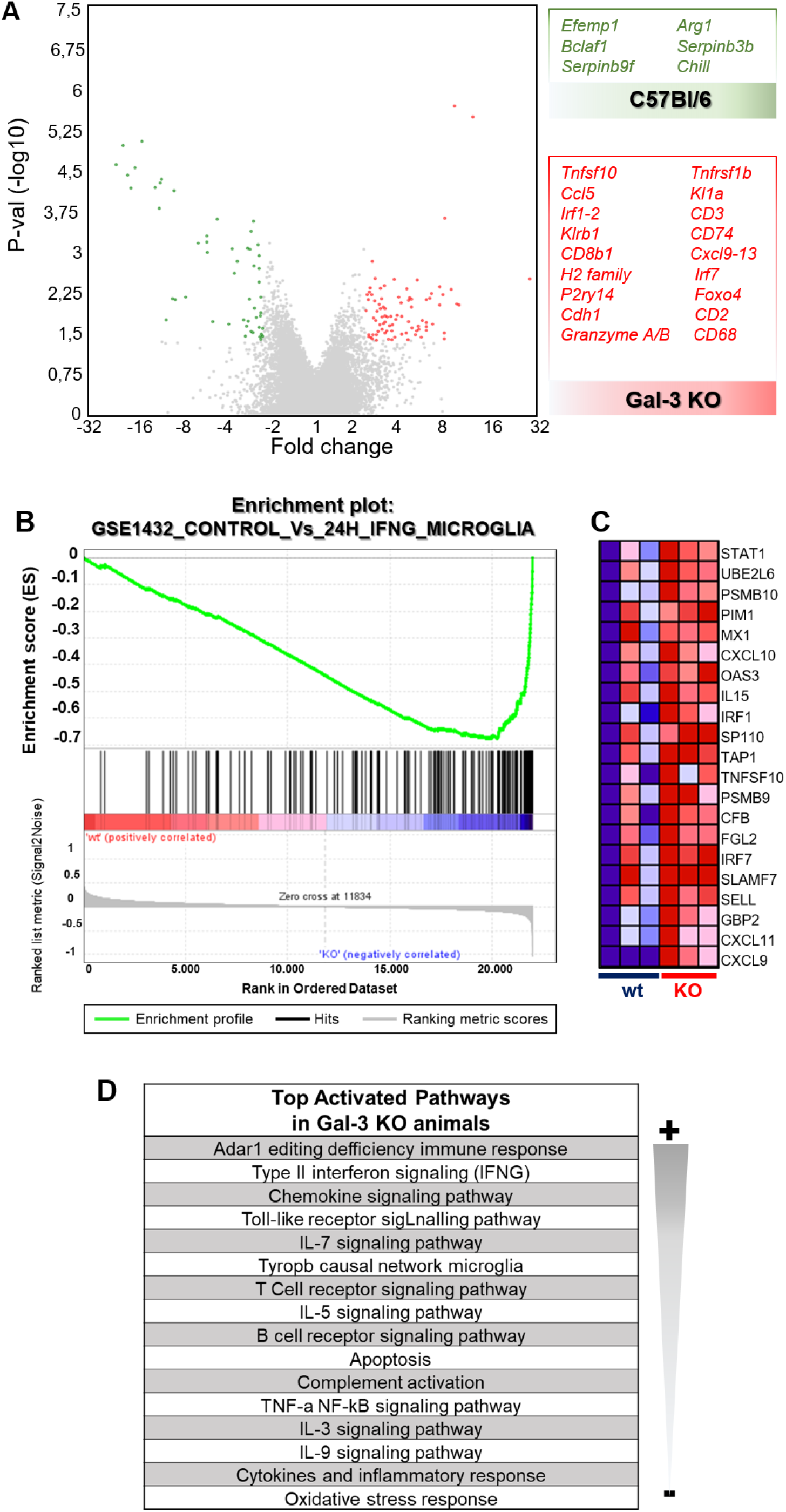
A.-) Volcano plot showing genes with significant upregulation (p<0,05 and <2-fold change) in wt (green dots) and Gal3KO mice (red dots). Next to the plot there are two tables with previously reported pro-metastatic genes (green), which are upregulated in wt mice, and pro-inflammatory/immune system-related genes (red), upregulated in the Gal-3 KO mice. B.-) GSEA plot showing the upregulation of genes involved in the IFNG response in microglia, which was upregulated in Gal3KO mice. C.-) Heatmap showing the 20 top dysregulated genes in the microglial IFNG gene set. D.-Most upregulated pathways in Gal3KO mice compared with wt.

### Glioblastoma

Comparing the TME of both experimental models, we observed 194 dysregulated genes, 111 were downregulated and 83 upregulated. Again, most dysregulated genes are typically expressed in activated immune system cells including microglia and macrophages (Clec7a, CD68, CD180) (Figure 8A). Again, GSEA showed the interferon gamma microglial pathway as one of the most upregulated clusters in the immunologic signature gene sets (Figure 8B)

**Figure 8.**
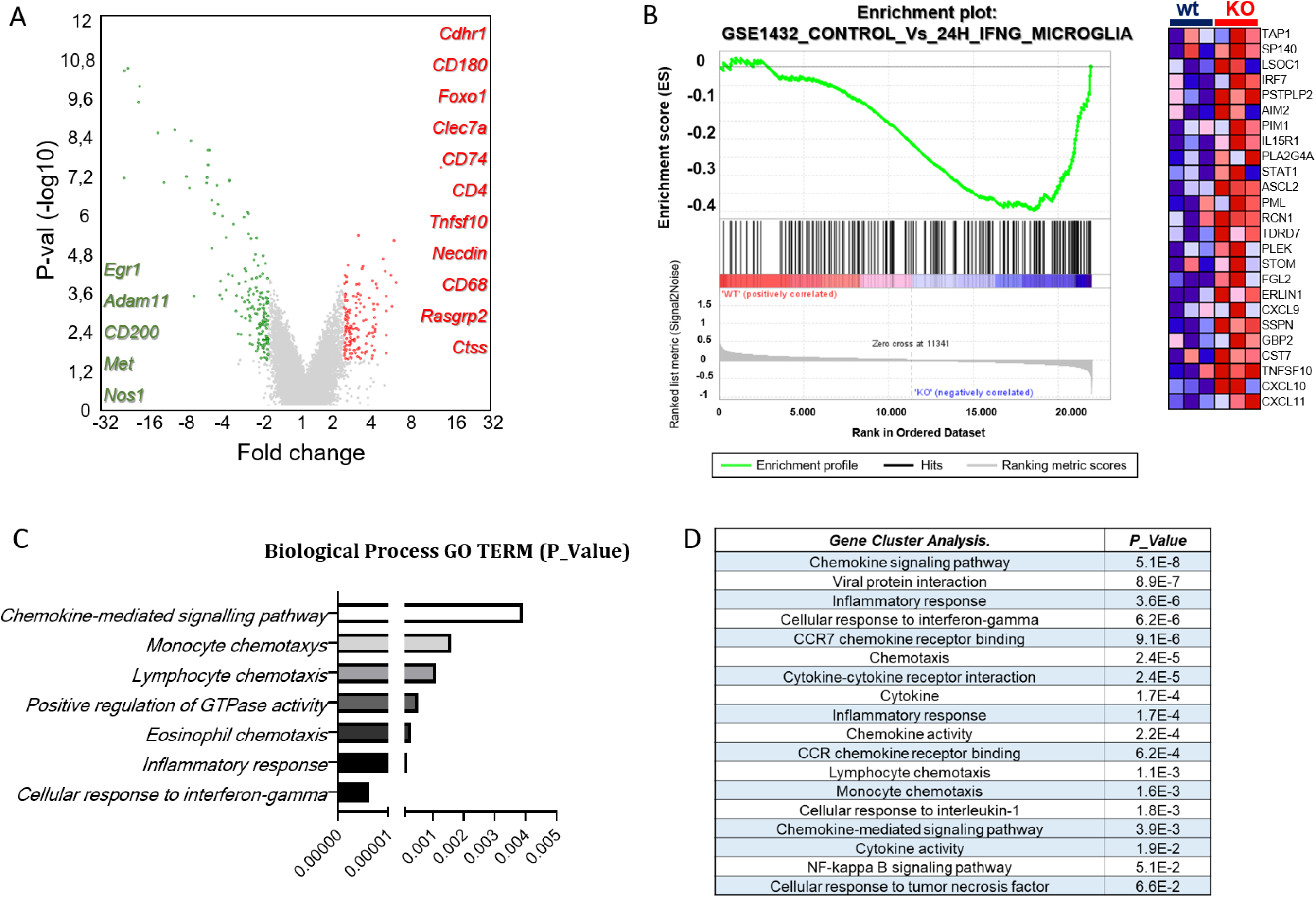
A.-) Volcano plot showing genes with significant upregulation (p<0,05 and <2-fold change) in wt (green dots) and Gal3KO mice (red dots). There are two columns depicting genes with previously reported pro-tumoural features (green), which are upregulated in wt mice, and pro-inflammatory/immune system-related genes (red), upregulated in the Gal-3 KO mice. B.-) GSEA plot showing the upregulation of genes involved in the IFNG response in microglia, which was upregulated in Gal3KOs after using TAC software. Gene cluster analysis after using DAVID software, showing the most dysregulated pathways.

Using the Database for Annotation, Visualization and Integrated Discovery (DAVID) we observed that GO TERM biological processes showed chemokine-mediated pathways, together with other myeloid-derived cells pathways clearly upregulated in Gal-3 KO mice, including again interferon gamma and inflammatory responses (Figure 8C). Finally, gene cluster analysis further validate our previous findings, with inflammatory-related pathways affected, such as Viral protein interaction, which is closely related with ADAR-1 (37), and cellular response to interferon gamma (Figure 8D).

Altogether, our transcriptomic analysis of the TME clearly show an immune-related pro-inflammatory microenvironment, which significantly affected BCBM and GB growth.

## 4. DISCUSSION

Galectin-3 is a member of the evolutionary conserved animal lectins family. It is expressed in several cells types playing key roles in different biological processes such as apoptosis, autophagy, inflammation, and cancer (9,38–40). However, in the last years, our group has unmasked the pivotal role of Gal-3 in immune system cells activation, specifically microglia (7,12,41). Thus, this work describes a novel role of this pleiotropic protein, activating an immunosuppressive phenotype in microglia in a brain tumour context.

We have observed how Gal-3 induces anti-inflammatory microglial activation by assessing levels of classical pro– and anti-inflammatory markers, in primary and immortalised microglia cell lines. Owing to the great plasticity of these cells, we found a complex behaviour, underlining the activating effect that exogenous addition of Gal-3 exerted on microglia. We found a strong upregulation of the classical immunosuppressive markers Arg-1 and PD-L1, mirroring the effects observed in microglia after the exposition to tumour-conditioned medium. Interestingly, such Gal-3-dependent anti-inflammatory response was abrogated once Gal-3 was inhibited. These results are in agreement with other works where blocking Gal-3 diminished levels of classical immunosuppressive markers (26,42). However, our *in vitro* results also suggest that tumoural Gal-3 plays an important role driving such immunosuppressive microglial response. In particular, the influence of Gal-3 in microglial PD-L1 expression is an appealing result since anti-PD-L1 therapies have experienced a great momentum in the clinic, although their results have not offered yet a significant increase in patients’ survival rates (28,43). Concomitant target of Gal-3 together with anti-PD-L1 therapies, may offer an alternative option to overcome the pro-tumoural expression of this immune checkpoint inhibitor within the TME.

Furthermore, Gal-3 increases migration and invasion in different tumour types (10,27). In agreement with this, we have found how the use of GB1107 (a specific Gal-3 inhibitor) dropped migration speed in breast cancer and glioblastoma cells. These results are in accordance with previous works (in tumours outside the brain), where reducing the levels of Gal-3 showed a direct correlation with cancer cells motility and invasion reduction (44,45), although no specific dependence of microglial Gal-3 had ever been established. We show how modulating microglial Gal-3 alters invasion properties of cancer cells, unveiling the promoter role of Gal-3 in two different types of brain-invading cancer cells.

We further pursued the interesting phenotypic shift found in microglia after Gal-3 modulation. There are numerous pre-clinical works describing the important role of Gal-3 promoting tumour growth and metastasis, predominantly in sites outside the central nervous system (10,44). However, its role in brain tumours has been less studied. Using a full Gal-3 knockout transgenic mouse, we observed a significant drop of the tumour burden in BCBM and GB syngeneic models. Despite BCBM and GB are a heterogeneous group of completely different diseases, we found a common pattern in both models: the TME in Gal-3 KO mice was shifted towards a pro-inflammatory state. This striking response was influenced by the fact that TAMs population showed jointly a decrease in anti-inflammatory markers together with a significant upregulation of pro-inflammatory enzymes, hence, supporting their tumouricidal immune response. It has been well described the important role of microglial Gal-3 in diverse neurological diseases (12,15). However, activation of microglia is regarded as a double-edged sword, and the changes in either pro-inflammatory or anti-inflammatory effects mediated by Gal-3 depend on the disease types, stages and severity (46). Thus, our study suggests that the immunosuppressive TME found in BCBM and GB could be partially reversed by increasing the population of iNOS-expressing TAMs after Gal-3 inhibition. Further supporting this hypothesis, we also observed how in both tumour models, the expression of the gold-standard immunosuppressive marker Arg1 (34) was significantly reduced in TAMs in Gal-3 KO mice.

Our transcriptomic analysis of the metastatic microenvironment showed a clear upregulation of key important genes involved in pro-inflammatory (anti-tumoural) immune responses such as histocompatibility complex II, antigen presentation and, importantly, T cells. These transcriptomic results were in agreement with our histological findings regarding the anti-tumoural CD8^+^/CD4^+^ TILs ratio. Furthermore, interferon and ADAR-1 pathways were also pathways clearly upregulated. Very recently, several studies have addressed the importance of ADAR-1 in promoting the immune response against tumours (37,47), highlighting the potential reach of our findings. All these results support the idea of Gal-3 inhibition as a key event to alter the fluctuating balance of pro– vs anti-inflammation within the TME, shifting TAMs towards a more pro-inflammatory state, favouring an anti-metastatic response.

As previously described, the transcriptomic analysis of the GB microenvironment also showed a clear upregulation of important inflammatory pathways in Gal-3 KO mice. It is noteworthy to emphasise the fact that our research group previously established TREM2, primarily expressed in microglia, as a pivotal ligand of Gal-3 (24). Recent studies have underscored the potential of TREM2 inhibition in enhancing anti-tumour activity within myeloid cells (35,48,49), thereby amplifying the efficacy of immunotherapies. In addition, the important role of RAGE as another ligand for Gal-3 has been recently described, thereby promoting glioma growth and invasion (50). In contrast to the inhibition of TREM2 or RAGE, our findings reveal that exclusive Gal-3 inhibition yields similar effects in restraining glioma growth. This is achieved by promoting a pro-inflammatory response in TAMs and supporting anti– tumoural gene pathways, which involve interferon-gamma, chemokines, and responses resembling inflammation.

In conclusion, this study furnishes compelling evidence highlighting the central role of Gal-3 in driving the polarisation of TAMs towards an immunosuppressive phenotype. This pro-tumoural state can be effectively reverted upon Gal-3 inhibition. Apart from the fact that Galectin inhibition has unquestionably emerged as a promising avenue in cancer research (51), it is worth noting that clinical trials targeting Gal-3 have been conducted for various diseases, albeit primarily outside the central nervous system, such as in COVID (NCT04473053), fibrosis (NCT01899859), or psoriasis (NCT02407041). Notably, for the first time, Gal-3 inhibitors are being explored for the treatment of central nervous system disorders (Alzheimeŕs disease-NCT05959239-). Consequently, Gal-3 inhibition emerges as a novel and promising pharmacological strategy for enhancing the immune response in patients afflicted with primary or secondary brain tumours

## Acknowledgments

The authors thank Dr Alberto Pascual for his assistance with transcriptomic software GSEA, and Dr T. Deierborg for providing the soluble Gal3 protein. We would also like to thank the “Genomic and Sequencing” services at the IBiS (Seville) and Dr Vasiliki Economopoulos for her in-house script to perform the colocalisation studies.

This work was supported by the European Union thanks to a Marie-Sklodowska Curie Action-IF (MSS, 795695), the Spanish FEDER I+D+i-USE (MSS, US-1264152), Asociación Española de Cáncer de Mama Metastásico (METAPREMIO 21, MSS) and the Ministerio de Ciencia e Innovación (MSS, PID2021-126090OA-I00).

## Authors Contributions

LCH, MTSM, ARR, JGR and MMA contributed to investigation, methodology and formal analysis. RT contributed to supervision, validation and investigation. JLV and MSS contributed to supervision, funding acquisition, administration and writing.

