## Supplementary material for "Galectin-3 depletion tames pro-tumoural microglia and restrains cancer cells growth": Supp Data

***ABSTRACT***

*The glycoprotein Galectin-3 (Gal-3) is a multifunctional molecule that plays a pivotal role in the initiation and progression of various central nervous system diseases, including cancer. Although the involvement of Gal-3 in tumour progression, resistance to treatment and immunosuppression has long been studied in different cancer types, mainly outside the central nervous system, its elevated expression in myeloid and glial cells underscores its profound impact on the brain's immune response. In this context, microglia and infiltrating macrophages, the predominant non-cancerous cells within the tumour microenvironment, assume critical roles in establishing an immunosuppressive milieu in diverse brain tumours. Through the utilisation of primary cell cultures and immortalised microglial cell lines, we have elucidated the central role of Gal-3 in promoting cancer cell migration, invasion, and an immunosuppressive microglial phenotypic activation. Furthermore, employing two distinct in vivo models encompassing primary (glioblastoma) and secondary brain tumours (breast cancer brain metastasis), our histologic and transcriptomic analysis show that Gal-3 depletion triggers a robust pro-inflammatory response within the tumour microenvironment, notably based on interferon-related pathways. Interestingly, this response is prominently observed in tumour-associated microglia and macrophages (TAMs), resulting in the suppression of cancer cells growth.*

SUPPLEMMENTARY DATA

### Cell culture and transfection

The murine glioma GL261, murine microglia BV2 and murine mammary carcinoma EO771 cell lines were purchased from the American Type Culture Collection. GL261 and BV2 were maintained in Dulbecco's Modified Eagle Medium (DMEM) (Gibco, USA) supplemented with 10% fetal bovine serum (FBS), 100 U/ml penicillin and 100 U/ml streptomycin (Invitrogen, USA). EO771 were maintained in RPMI 1640 (Gibco) supplemented with 10% FBS, 100 U/ml penicillin and 100 U/ml streptomycin (Invitrogen). Cells were regularly tested for mycoplasma contamination using the Mycoblue Mycoplasma Detector Kit (Neo Biotech ref. #NB-54-0011). Cell lines were maintained in a humidified 5% $CO_2$/95% air environment at 37°C.

Cells were knocked down by small interfering RNA (siRNA) targeting galectin-3 (Santa Cruz Biotechnology, USA ref: sc-155994-SH) according to the manufacturer's siRNA transfection protocol. Briefly, galectin-3 shRNA or scramble plasmids were mixed with shRNA transfection reagent (sc-108061) and shRNA transfection media (sc-108062). Then, the plasmid mixture was transferred into the cells after 6h incubation. Cells were then cultured in DMEM supplemented with 20% FBS, 100 U/ml penicillin and 100 µg/ml streptomycin. After 48 h, transfected cells were selected using 1 µg/ml of puromycin in normal culture media. Non-transfected cells were removed, and transfected cells were maintained for verification using qPCR and Western blotting.

**Invasion and migration studies**

BV2, GL261 and EO771 cells were co-cultured using transwell clear inserts, polyester (PET) membrane (Corning, USA ref: 3470). In vitro cell invasion for these cell lines was assessed after 24h using a corning BioCoat matrigel invasion chamber assay (Corning, ref: 354480). The experiment and subsequent measurement of cell invasion was performed according to the manufacturer's Cell Invasion Assay protocol. Briefly, the cell lines were incubated in Corning BioCoat Matrigel Invasion Chambers for 24h in a humidified tissue culture incubator at 37°C, 5% CO2. After incubation, non-invading cells were removed from the top of the membrane by scrubbing, and cells on the bottom of the membrane were stained to allow visualisation.

Cell migration studies were performed using the scratch wound assay method. Cells plated on a chamber slide were at appropriate confluence, and a 600µm-wide wound was scratched through the centre of the well using a 200 µl pipette tip. Cells were placed on the microscope and images were taken at 6 and 24h after scratching (Supplementary Figure 1A).

Recombinant galectin-3 (Gal-3) was produced by the Lund-Protein Production Platform (Lund University, Sweden). Briefly, Gal-3 production was performed in strain E. coli TUNER(DE3)/pET3c-hum-Gal3 grown in LB medium, 18 °C, 250 rpm with 1 mM IPTG overnight. After cell lysis and ultracentrifugation, Gal3 was purified on a 20-ml lactocyl-sepharose column. Peak fractions containing Gal3 were pooled and dialyzed against phosphate buffer saline (PBS, Nzytech, Lisbon, Portugal), pH 7.4.(15)

**RNA extraction and quantitative PCR**

RNA was extracted from the cell lines using the RNeasy Kit (Qiagen, Netherlands). Subsequently, 0.5 µg of total RNA was converted into cDNA using the First-Strand cDNA Synthesis Kit (Thermo Fisher Scientific, USA). The indicated primers (Table 1) were used for qPCR and performed on the LightCycler 480 (Roche Molecular Systems Spain). Each experimental group was performed in triplicate to obtain the mean cycle time (CT).

Table 1. List of qPCR primers

| **Genes** | **Sense** | **Primers (5'-3')** |
|---|---|---|
| *Nos2* | Forward | GAGCCACAGTCCTCTTTGC |
| | Reverse | CTCTCTTGCGGACCATCTCC |
| *Arg1* | Forward | GTTGATGTCCCTAATGACAGC |
| | Reverse | CATTCTTCTGGACCTCTGCC |
| *Tnf* | Forward | TGCCTATGTCTCAGCCTCTTC |
| | Reverse | GAGGCCATTTGGGAACTTCT |
| *Pdcd1* | Forward | CTACGGGCGTTTACTATCACGG |
| | Reverse | AGGGAATCTGCACTCCATCG |
| *Lgals3* | Forward | GATCACAATCATGGGCACAG |

| | | |
|---|---|---|
| | Reverse | GTGGAAGGCAACATCATTCC |

**Western blotting assay**

Total proteins were harvested using lysis buffer (Cell Signaling Technology, USA) and stored at 4°C for 30 minutes. The protein content of the samples was estimated by the micro-Lowry method using bovine serum albumin as a standard, with 25 μg of protein loaded for each lane. Protein samples were separated by SDS-PAGE (10%) and transferred to a nitrocellulose membrane (Novex; Life Technologies, USA). Membranes were blocked with blocking buffer (5% milk in TBS: 20 mM Tris-HCl, pH 7.5, 500 mM NaCl and 0.05% Tween 20) for 1 hour at room temperature. Membranes were then incubated with anti-Gal 3 antibody (1:1000). GAPDH antibody (Sigma-Aldrich; 1:2000; USA) was used as a loading control.

**Animals**

Experiments were performed in 12-week-old mice C57BL/6 and galectin-3 null mutant mice on the same background, both lines obtained from Charles Rivers. Male mice were used for the GB model and female mice for the BCBM model. Animal experimentation was carried out in accordance with the European Community Council Directives (86/609/EU) and Spanish law (R.D. 53/2013 BOE 34/11370-420, 2013) for the use and care of laboratory animals. All the animal procedures in this study were previously approved by ethics committee of University of Seville. Animals were housed under a 12h light/dark cycle with free access to food and water.

**Surgery**

Female C57BL/6 mice were injected with 1μl of PBS containing 103 syngeneic mouse mammary carcinoma EO771 cells by unilateral stereotaxic intracerebral injection (in the left striatum) at the following coordinates: +0.5 mm/+1.9 mm lateral to bregma, 2.8 mm deep, over a period of 5 min. Using a fine glass microcapillary (70 μm tip).

Male C57BL/6 mice followed the same surgical protocol as above, but instead, 1μl with 104 syngeneic mouse glioblastoma cells GL261 were stereotactically injected.

**Immunohistochemistry**

Coronal sections (10 μm thick) of brains were cut using a cryostat (Leyka, Germany) at -20°C, mounted in Superfrost slides (Fisher Scientific USA) and allowed to dry at room temperature (RT) overnight. Brain sections were then frozen at -20°C until immunohistochemistry was performed. Different antibodies were used against Iba-1 (rabbit anti-Iba1 1:1000, Wako ref; 019-19741) mouse anti-Gal3 (1:1000) (Sigma Aldrich ref: 255M-18), goat anti Arg1 1:100 (Santa Cruz ref: sc18351), rabbit anti iNOS 1:150 (Abcam UK ref: ab3523), rat anti CD80 (Invitrogen ref:MA1-81972), rabbit anti CD206 1:200 (Cell signalling ref:24595) and rabbit anti PD-L1 1:200 (Cell signalling ref: 86744).

The sections were washed in 0.1% PBS-Tween. Antigen retrieval was then carried out with citrate buffer 20 min at 90°C. They were then left at RT for 15 minutes. The sections were then blocked with 5% BSA in PBS for 30 minutes and then incubated overnight at 4°C with the appropriate primary antibodies. The next day, brain sections were tempered at RT for 20 minutes and washed in PBS. They were then incubated with the appropriate secondary antibodies: Donkey anti Mouse Alexa 488 (Life technologies USA ref: A21202), Donkey anti Rabbit Alexa 647 (Life technologies ref: A31573), Donkey anti Goat Alexa 488 and 647 (Life Technologies ref: A11055, A21447), Chicken anti rat Alexa 594 (Life Technologies ref: A21471) for 45 minutes at RT. The sections were then washed and stained with DAPI solution for 5 minutes and mounted on coverslips.

**Confocal microscopy**

Sections were imaged using a Leica Stellaris 8 STED confocal laser scanning microscope with a ×20 objective. Images of wild-type and transgenic mice were taken on the same day and under the same conditions (laser intensities and photomultiplier voltages). Morphometric analysis of fluorescence-labelled structures was performed offline using ZEN software and Fiji ImageJ (W. Rasband, National Institutes of Health).

**Quantitation of tumour burden**

All data were analysed blinded to experimental conditions. To assess tumour growth, as previously described, 10 µm thick sections were cut across the entire striatum. All data were analysed blind to experimental conditions. ImageJ® was used to quantify tumour area for each animal (n = 6 per group) on at least 8 sections evenly distributed through the striatum, from the beginning to the end of the tumour. Tumours were circumscribed and compared to the total brain area analysed (16). Tumours were circumscribed and compared with the total brain area analysed (Supplementary Figure 2)

**TAMs analysis**

Immunofluorescence images were captured using an inverted confocal microscope Leica Stellaris 8 STED. Images were analysed using the ImageJ software package (NIH; ref. 23). Full information on the colocalisation studies can be found in the Supplementary Data.

Images were analysed using in ImageJ software package (NIH). A custom-written code was designed to calculate the Pearson's correlation coefficient, the Mander's overlap coefficient, and the co-localisation coefficients m1 and m2, before quantifying co-localisation between two fluorescent markers. This code enabled the percentage of co-localisation, as well as the percentage of area in the image of each marker that was above the user-defined threshold, to be determined. The signal arising from targeted proteins were normalised to microglia area and expressed as density of m1 pixels per m2 area. For all animals, images were taken at the same time and under the same conditions. A second code was created to calculate the percentage of co-localising pixels, using the same algorithm but modified for three fluorescent markers. The degree of co-localisation between iNOS/Arg1 and Iba1 was quantified by counting the number of green pixels within the red channel using the first code.

The relative contribution of microglia Arg1 and iNOS expression was calculated from the number of positive pixels and percentage of co-localisation according to:

Total{npixels(iNOS/Arg1)}={npixels(Iba1)×%colocalisation(iNOS/Arg1/Iba1)}

In this equation, total{npixels(iNOS/Arg1)} corresponds to the number of pixels in microglia and %colocalisation to the percentage of co-localisation of iNOS/Arg1 per Iba1-positive pixels.

**Microarray analysis**

To study the differences in genes expression between wt and Gal-3 KO mice, we used Transcriptome Analysis Console (TAC) Software and selected genes that had at least a 2-fold change in expression and a t-test P-value of less than or equal to 0.05. We also used Gene Set Enrichment analysis (GSEA) and the Database for Annotation, Visualization and Integrated Discovery (DAVID) to further analyse gene clusters comparisons and key biological pathways.

The RNA quality was analysed using Agilent 2100 Bioanalyzer (Agilent). Samples with RNA integrity number (RIN) below 7 were pdiscarded. RNA was amplified and labelled using the GeneChip WT Pico Reagent Kit (the total RNA isolated was used as the starting material; Affymetrix). The amplified cDNA was quantified, fragmented, and labelled in preparation for hybridisation with the mouse Clariom S Assays (Thermo-Fischer). C57Bl/6 (WT) and Gal-3 KO (Gal3KO) tumour samples were subjected to this analysis. One hundred ng of RNA was used for the subsequent labelling reaction. cRNA and single strand (ss) cDNA were synthesised using Affymetrix GeneChip WT Plus Reagent according to the manufacturer's instructions. The ssDNA (5.5 μg) was then fragmented and biotin-labelled using the GeneChip WT Terminal Labeling Kit (Thermo-Fischer) according to the manufacturer's instructions. The labelled cRNA was hybridised to the microarray for 17h using the GeneChip Hybridisation, Wash and Stain Kit (Thermo-Fischer). To visualise fluorescence signals, the microarray was scanned using the GeneChip Scanner 3000 7G.

To identify underlying biological processes in endothelial cells from Gal-3KO mouse models, we used GSEA. We analysed the enrichment of 5219 gene sets from the Immunologic Signature Database C7.

**Statistical analysis**

All individual measurements constitute independent biological replicates and the experiments were repeated at least three times (for quantifications, the n is specified in the figure legends). All data are presented as mean ± SD. Unless otherwise stated, data were analysed using GraphPad Prism (v8.4). For 2 groups, Student's t-test (2-tailed) was used. For more than 2 groups, 1-way ANOVA with Tukey's multiple comparison test or two-way Anova with Bonferroni test were used and adjusted P values are reported. Statistical significance was defined as $p < 0.05$.

SUPPLEMENTARY FIGURES

**SUPP FIG 1**

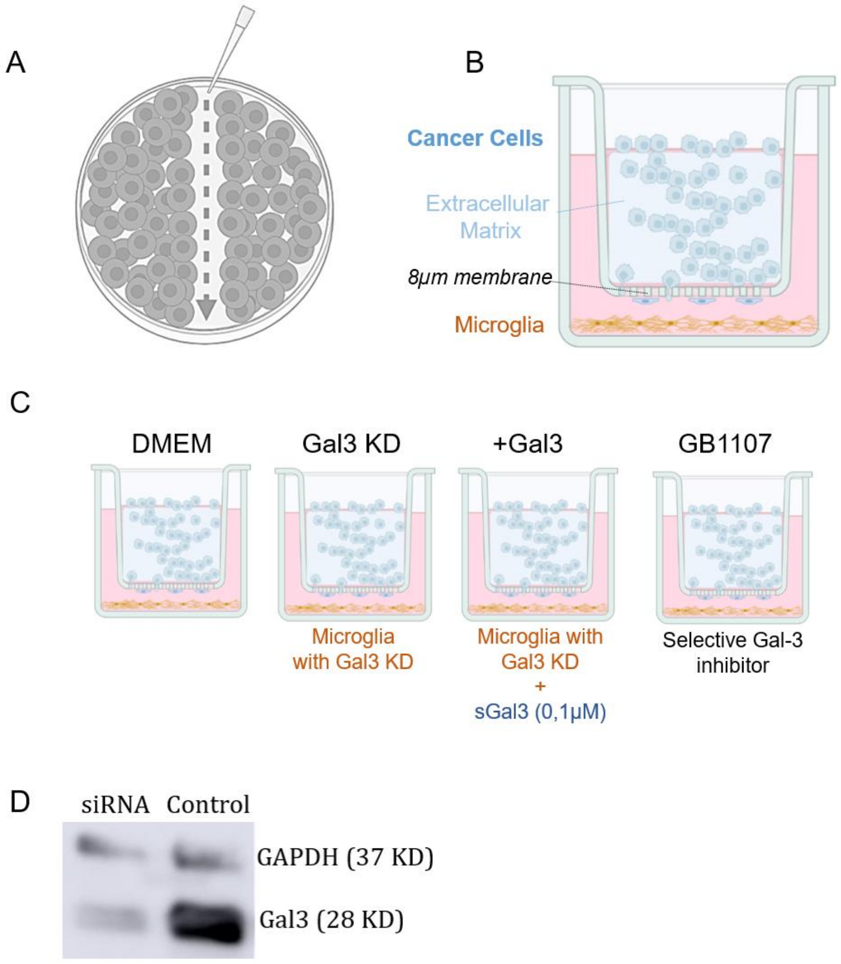

**Supp Fig 1**. A.- Scheme of the 2D scratch assay performed with EO771 and GL261 cells. B.-) Schematic of the invassion assay with the different cell components. C.- The four experimental approaches during the invasion assay are depicted as DMEM (control conditions), GAL3KD (siRNA), +Gal3 (0,1µM) and the use of GB1107 in parental BV2 conditions. D.-) Western blot showing Gal-3 KD level in BV2 cells.

**SUPP FIG.2**

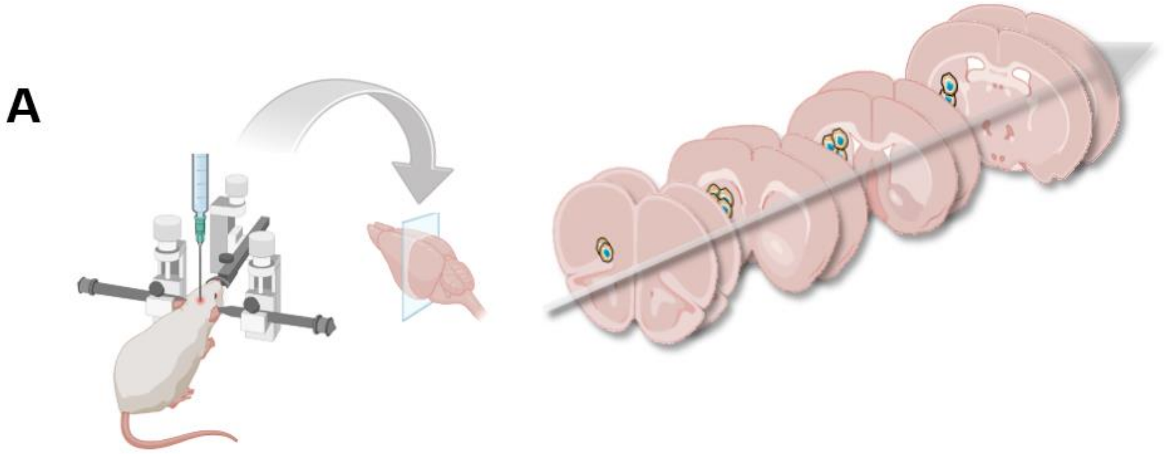


**Supp Fig 2**. Schematic of the quantitation performed throughout the whole striatum of the mice. 10µm-thick sections were collected and tumour area quantified from the beginning to the end of the tumour. At least 8 sections were used to perform the tumour burden measurement.

**SUPP FIG. 3**

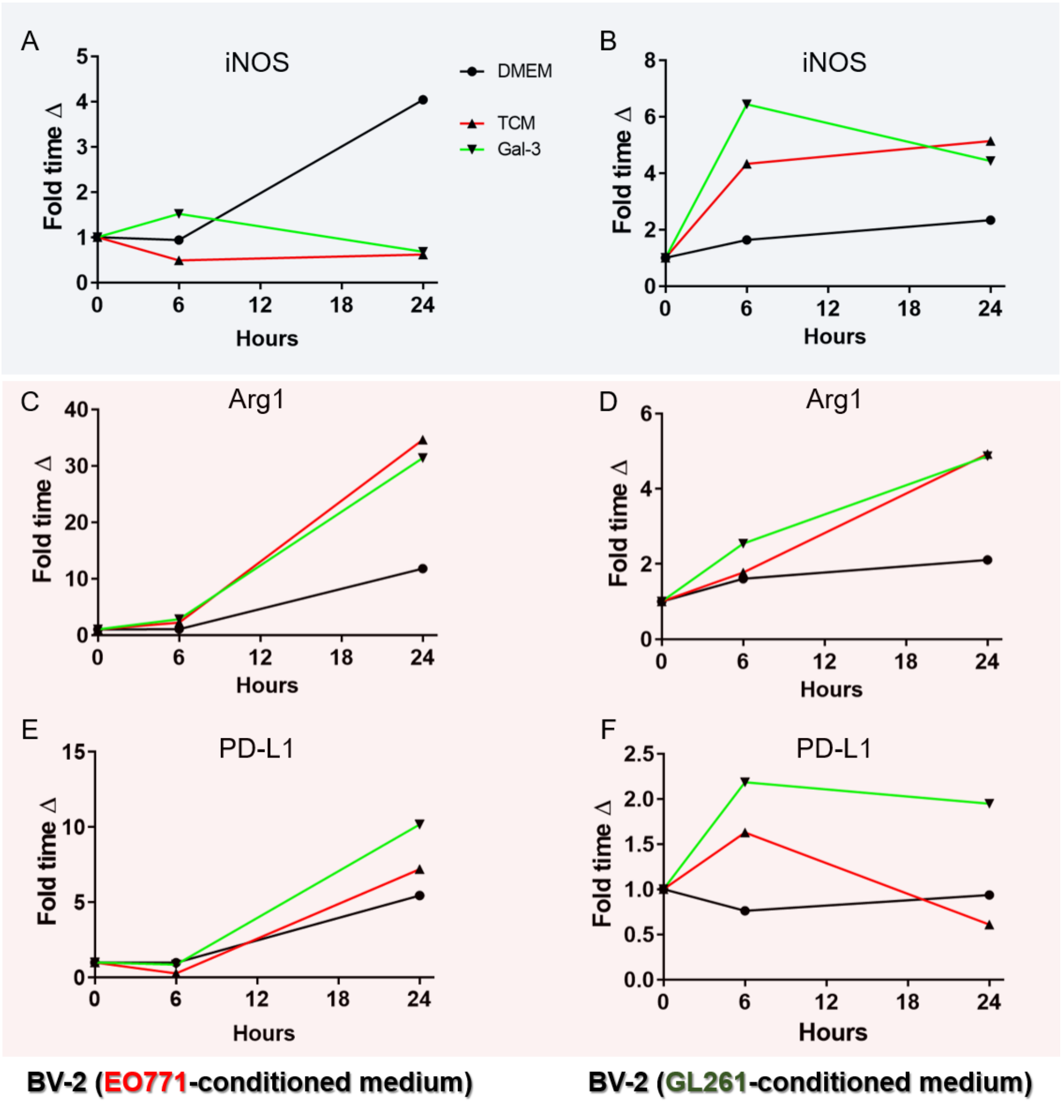


**Supp. Fig 3**. Graphs showing qPCR results of the expression of pro-inflammatory iNOS in BV-2 cells coated with EO771-conditioned medium (left column) and GL261-conditioned medium (right column). The same analysis for Arg1 (C-D) and PD-L1 (E-F). Every markers was analysed under control conditions (DMEM), TCM and exposition (24h) to exogenous Gal-3.

**SUPP. FIG 4**

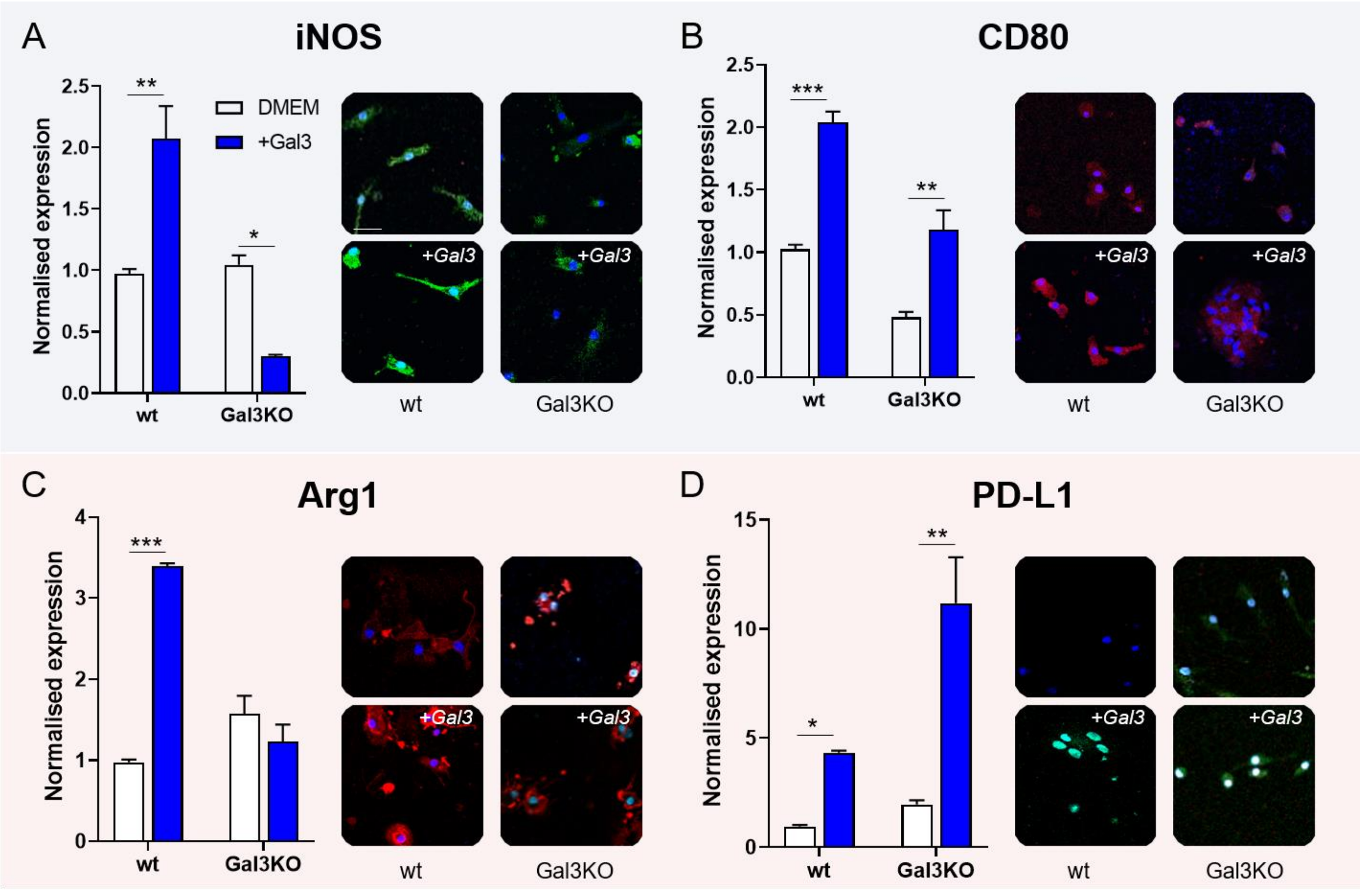


**Supp. Figure 4.** Primary microglia from wt and Gal-3 KO P1-3 mice. A.-) Graph showing comparison of iNOS expression in normal medium (control conditions, DMEM, White bars) and after the addition of exogenous Gal-3 (24h, 0,1µM, Blue bars). Pictures show expression of iNOS (green) and nuclei (DAPI) in microglia cultured in DMEM (top row) or Gal-3 (bottom row). B.-) As per A, measuring expression of pro-inflammatory CD80. C.-) Immunosuppressive Arginase1 marker measured as per A. D.-) Graphs and images depicted as per A. Two way anova and Bonferroni post hoc test. **, p<0,001, ***, p<0,005

**SUPP FIG 5**

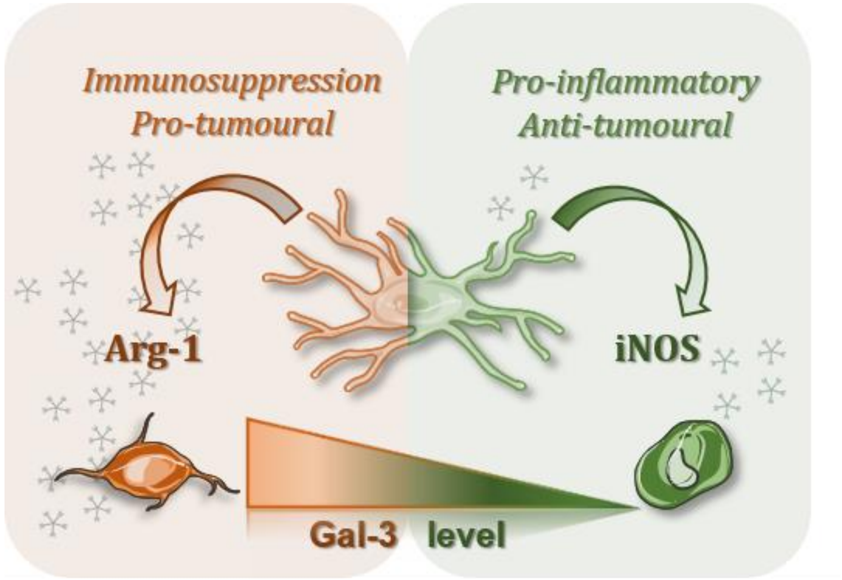


**Supp Fig 5**. Schematic of the role of Gal-3 in microglia. When the presence of Gal-3 is upregulated, microglia tend to overexpress arginase-1. Whilst, on the contrary, if Gal-3 levels are low, microglia show increase of the expression of iNOS. Galectin-3 is depicted as 5-pointed stars at the back.
